# About a tetraploid ivy in Sicily: from autochthonous *Hedera* to horticultural-invasive-hybrid package?

**DOI:** 10.1101/751743

**Authors:** Alain Fridlender, N. Pech

## Abstract

In Sicily, *H. helix* is the unique known native species whereas *H. helix* susbsp. *poetarum* is putatively naturalized in some forests and *H. canariensis* cultivated in various urban’s garden. Trichome morphology and genome size of some ivies from various west Mediterranean forests were compared to Sicilian ones. Ivies from southern Italy, continental France, Corsica and Mallorca belong to typical European diploid stellate trichomes *Hedera helix*. Hexaploid ivies from southern Spain have been identified as native *H. iberica*. Contrariwise, Sicilian ivies studied are related to western European *H. hibernica* (tetraploids with stellate trichomes). Is *H. helix* the most widespread and indigenous ivy in Sicily? Therefore, it would be the first time that tetraploid would be reported in Sicily where it could possibly correspond to an unnoticed autochthonous taxon. However, our results let us think it rather represents an invasive which impact on this island rich in endemic species could be considerable.

## 1. Introduction

Species belonging to Eurasian genus *Hedera* L. (Araliaceae) are notoriously difficult to identify due to high leaves polymorphism caused by environmental conditions and heteroblasty: ontogenic change from juvenile to adult leaves (Robbins 1957, 1960; Frydman et al. 1973a, 1973b). As heteroblastic development (partially under influence of the hormonal production from roots and buds) is reversible (transition from adult to juvenile), we may find a great variety of leaves shape within one individual and all along one stem and its branching.

According to authors, *Hedera* Eurasian genus would include 12 to 16 taxa (Lum and Maze 1989; McAllister and Rutherford 1990; Vargas et al. 1999; Ackerfield 2001). Thanks to trichome morphology - stellate versus scale-like-combined with ploidy levels, recent works allow us to better characterize them (Table 1).

**Table 1.** Ploidy level and trichome morphology of ivies (genus *Hedera* L., Araliaceae).

| Ploidy level | Diploid<br>2n = 48 | Tetraploid<br>2n = 96 | Hexaploid<br>2n = 144 | Octoploid<br>2n = 192 |
| --- | --- | --- | --- | --- |
| Stellate<br>trichomes | *H. azorica*<br>*H. helix* L. subsp. *helix*<br>*H. helix* subsp. *poetarum*<br>*H. helix* subsp.<br>*rhizomatifera*<br>*H. helix* subsp. *caucasigena* | *H. Hibernica* |  |  |
| Scale trichomes | *H. maroccana*<br>*H. rhombea*<br>*H. nepalensis*<br>*H. canariensis* | *H.*<br>*algeriensis* | *H. cypria*<br>*H.*<br>*pastuchovii*<br>*H.*<br>*maderensis*<br>*H. iberica* | *H.*<br>*colchica* |

The five easternmost species of the genus area distribution present scale like trichomes: *Hedera cypria* McAllister (Cyprus), *H. colchica* (K. Koch) K. Koch (South and Est Black Sea territories), *H. pastuchovii* Woronow (West and South Caspian Sea territories), H. *nepalensis* Koch. (Afghanistan to China), *H. rhombea* (Miq.) Bean. (Korean peninsula, Japan, Taiwan) (Figure 1a).

**Figure 1.**
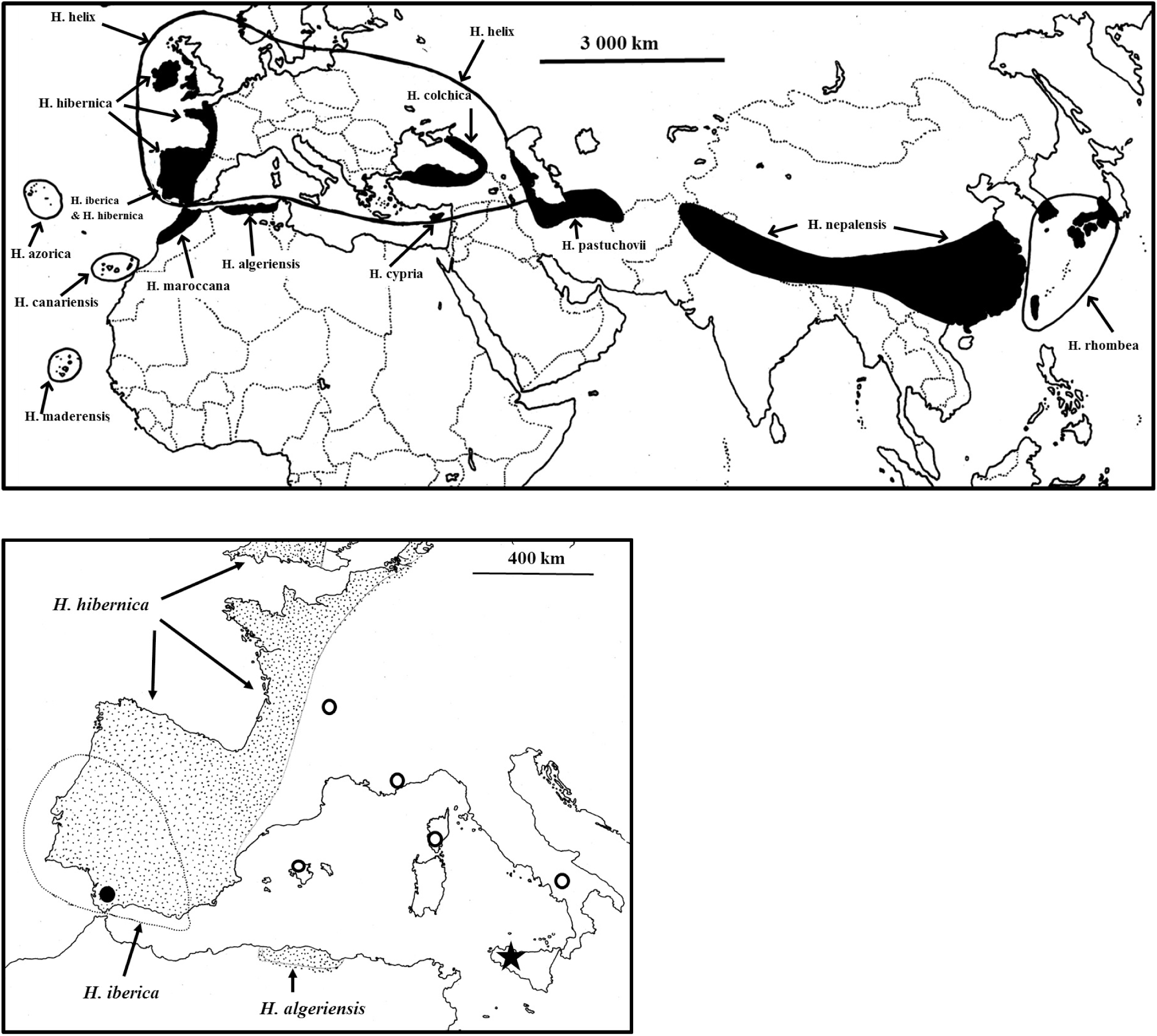
General outline of native distribution of ivies. **1a** - Distribution of the 13 more commonly accepted species of *Hedera*. **1b -** Distribution map of *H. iberica* (6x) and the two tetraploid ivies species (dotted): Atlantic western European *H. hibernica* and endemic north Algerian *H. algeriensis*. Studied populations: Southern Spain hexaploid population from Los Barrios (black dot); diploid populations from France, Italy, Corsica and Mallorca (withe dots); Sicilian tetraploid populations of Alcamo and Madonie (star).

*Hedera helix* L., a diploid stellate trichome plant is present in all European countries: within numerous infraspecific taxa described, the most noticed are *H. helix* subsp. *rhizomatifera* McAllister in southern Spain and *H. helix* subsp. *caucasigena* Kleop. in Turkey and Caucasus. Native *H. helix* s.l. distribution remains unclear in particular where several species and infraspecific taxa live in sympatry. In Ireland and in Western part of Great Britain, France and Iberian Peninsula grows *H. hibernica* (G. Kirchner) Bean. *H. helix* s.l. and *H. hibernica* are invasive species in North America, Argentine, Chile, Australia, New Zealand…

South westernmost part of *Hedera* distribution presents the highest ivies diversity. Each one among the three Atlantic archipelagos possess endemic species: *H. canariensis* Willd., *H. maderensis* K. Koch ex A. Rutherf, *H. azorica* Carr. In North Africa, the genus is represented by two endemics species with scale-like trichomes (*H. maroccana* McAllister, *H. algeriensis* Hibberd). Five taxa are known in Iberian Peninsula: three stellate trichomes species (*H. helix* subsp. *helix, H. helix* subsp. *rhizomatifera, H. hibernica)* and two scale-like trichomes taxa (*H. iberica* (Mc Allister) Ackerfield & J. Wen [= *H. maderensis* K. Koch ex A. Rutherf. subsp. *iberica* Mc Allister], and locally naturalized *H. maroccana*).

In the field, it is not so easy to identify ivies! Leaves shape varies greatly according to environmental conditions (light, humidity), it is different on horizontally and vertically crawling shoot-youth leaves much more cut than adult ones. Finally, to observe trichomes, it is necessary to use a more powerful magnifying glass than the ones usually available on the ground. Trichomes are abundant in apex, more dispersed on young leaves and usually absent on mature leaves. Moreover, trichomes fall easily when touched (harvesting, herbarium handing…).

In addition, in Europe ivies are largely cultivated for a long time and offered for sale in nursery since the 18th century: *H. hibernica, H. maroccana, H. algeriensis* and *H. colchica* have now commonly escaped from garden.

Furthermore, numerous hybrid and cultivar ivies are introduced, sometimes escape from gardens and remain even more difficult to identify. Triploid clones (most of them sterile) apparently do not exist in native European population but they are sold in the markets and escape in the field. Triploids and particularly allotetraploids may appear spontaneously in areas were introduced populations come into contact with native ivies or other previously naturalized ivies cultivars. Hybrids individuals appear to be more and more common in Europe (Marshall et al. 2017).

Despite the huge diversity of phenotype and ploidy levels, botanists, naturalists or ecologists name generally the ivies indistinctly under the binomial *Hedera helix* L. In this study, we use cytometric analyses and trichomes morphology to 1) better identify ivies collected from western Mediterranean islands, 2) point out taxonomic inconsistencies and 3) highlight the importance of accurate species identification in a context of global change.

## 2. Material and methods

### 2.1. Plants studied

The origins and numbers of the plants for each of the studied population are shown in Table 2. Cytometric analyses were performed on one Iberian population of *Hedera iberica* (Figure 2) and various *Hedera* cf. *helix* look like plants from three continental populations (Italy, France) and three insular origins: Mallorca, Corsica and Sicily (Figure 1b).

**Table 2.**
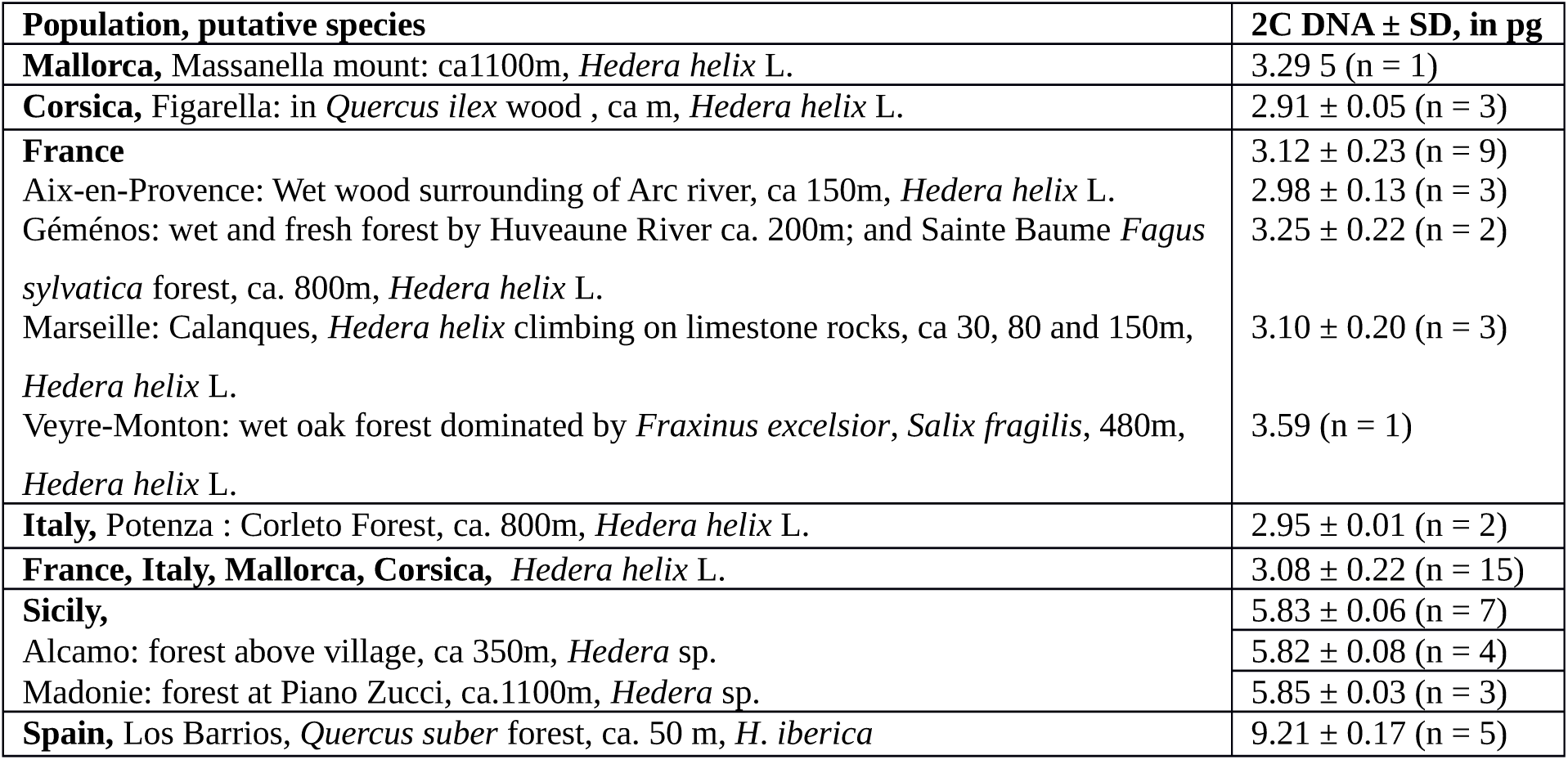
Origin of studied populations of *Hedera* and Genome size (2C DNA in picograms ± standard deviation). **n =** number of individuals measured. **Marseille:** plants were collected in the center of Massif des Calanques because on the edge of Calanques, ivies are a taxonomic mixture: most individuals are hybrid between native ivy and introduced cultivars, most of them issued from *H. algeriense* and “English ivy”.

**Figure 2.**
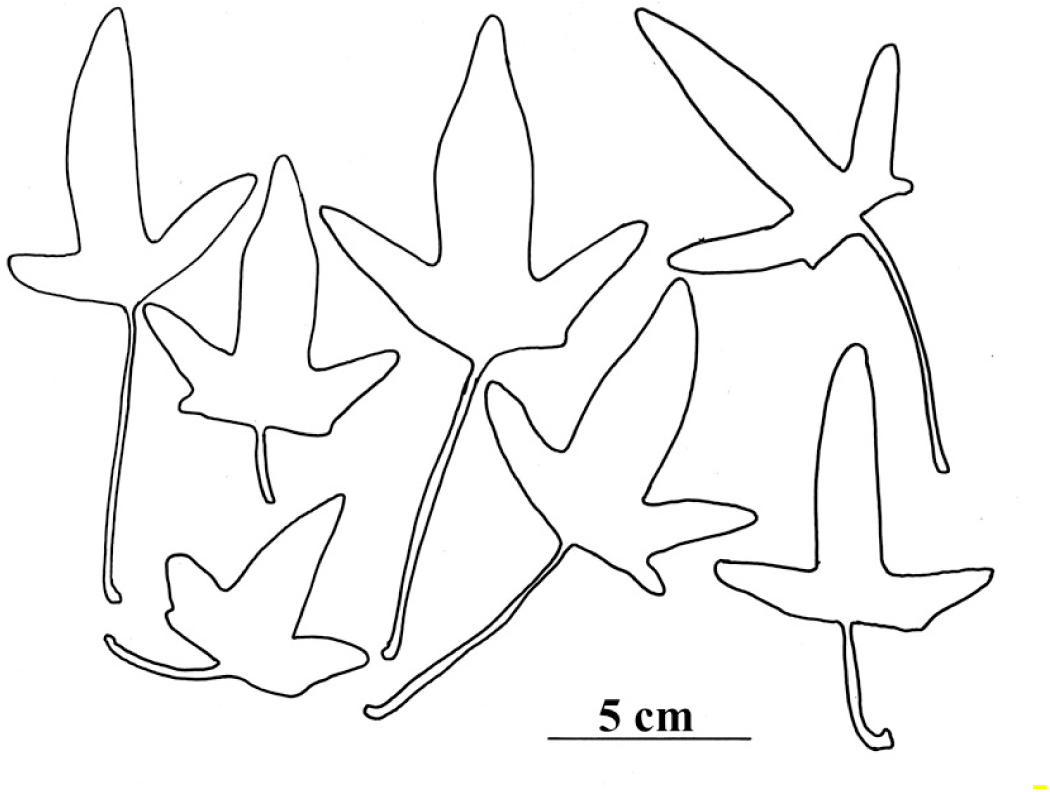
*Hedera iberica* leaves shape from los Barrios (Spain) population.

### 2.2. Genome size analysis by flow cytometry

The analyzed leaves were collected in the wild and immediately sent to the laboratory. The total nuclear DNA amount was assessed by flow cytometry (Marie and Brown 1993; Fridlender et al. 2002). For Iberian ivies, we used *Artemisia arborescens* L. from Crete (2C = 11.43 pg) as an internal standard, *Petunia hybrida* cv PxPc6 (2C = 2.85 pg) for Sicilian ivies and for all other plants *Lycopersicon esculentum* (2C = 2.01 pg). Leaves of the internal standard and *Hedera* were chopped using a razor blade in a plastic Petri dish with 600 μl of Galbraith nucleus-isolation buffer (Galbraith et al. 1983) containing 0.1% (w/v) Triton X– 100, supplemented with 10 mM sodium metabisulphite, 1% (w/v) polyvinylpyrrolidone 10,000 and RNAse (2.5 U/ml). The suspension was filtered through 50 μm nylon mesh. The nuclei were stained with 75 μg/ml propidium iodide, a specific DNA fluorochrome intercalating dye, and kept 15 min at 4°C. DNA content of 5,000–10,000 stained nuclei was determined for each sample using a cytometer (CyFlow SL3, Partec, Munster, Germany). The total 2C DNA value was calculated using the linear relationship between the fluorescent signals from stained nuclei of the *Hedera* individuals and the internal standard. The mean value was calculated from measurements of samples comprising 1 to various individuals according to populations (Table 2).

### 2.3. Statistical analysis

The relationship between ploidy P and 2C DNA content x for known ploidy level was modeled considering the following quadratic model: Pi=a + b xi + c xi^2^ + Ei. Ei terms are supposed to be identically and independently distributed according to a centered Gaussian distribution whose standard deviation is indicated by σ. Significance of parameters was tested using classical t-test. Statistical analyses were performed using R (R Core Team, 2019).

### 2.4. Trichomes

Trichomes from young leaves, apex and shoots from dried specimens were observed in SEM according to classical microscopical protocol. Measurements of various parameters (arms number, length of fused arms, lengh of free part of arms) from 25 trichomes from Sicily and 15 from Spain were done, based on Lume and Maze (1989).

## 3. Results

### 3.1. Trichomes morphology

Andalusian ivies have leaves shape with terminal leaf lobe much longer than the others (Figure 2) and scale-like trichomes (Figure 3a-b). Trichomes from young dried leaves from Los Barrios ivy population have 8-14[17] arms (m = 9.2 ± 1.5) fused in a 48-66 µm central disc diameter (m = 55.2 ± 7.8). Arms free length part varies from 55 to 200 µm (m = 118.7 ± 40.7) with basal width of 14-27 µm (m = 20.4 ± 3.49). Plants do not present difficulty of identification based on trichomes morphology (Ackerfield & Wen 2002) and limbs shape (Valcȧrcel et al. 2002): all the individuals correspond to endemic *H. iberica* (McAllister) Ackerfield & J. Wen.

**Figure 3.**
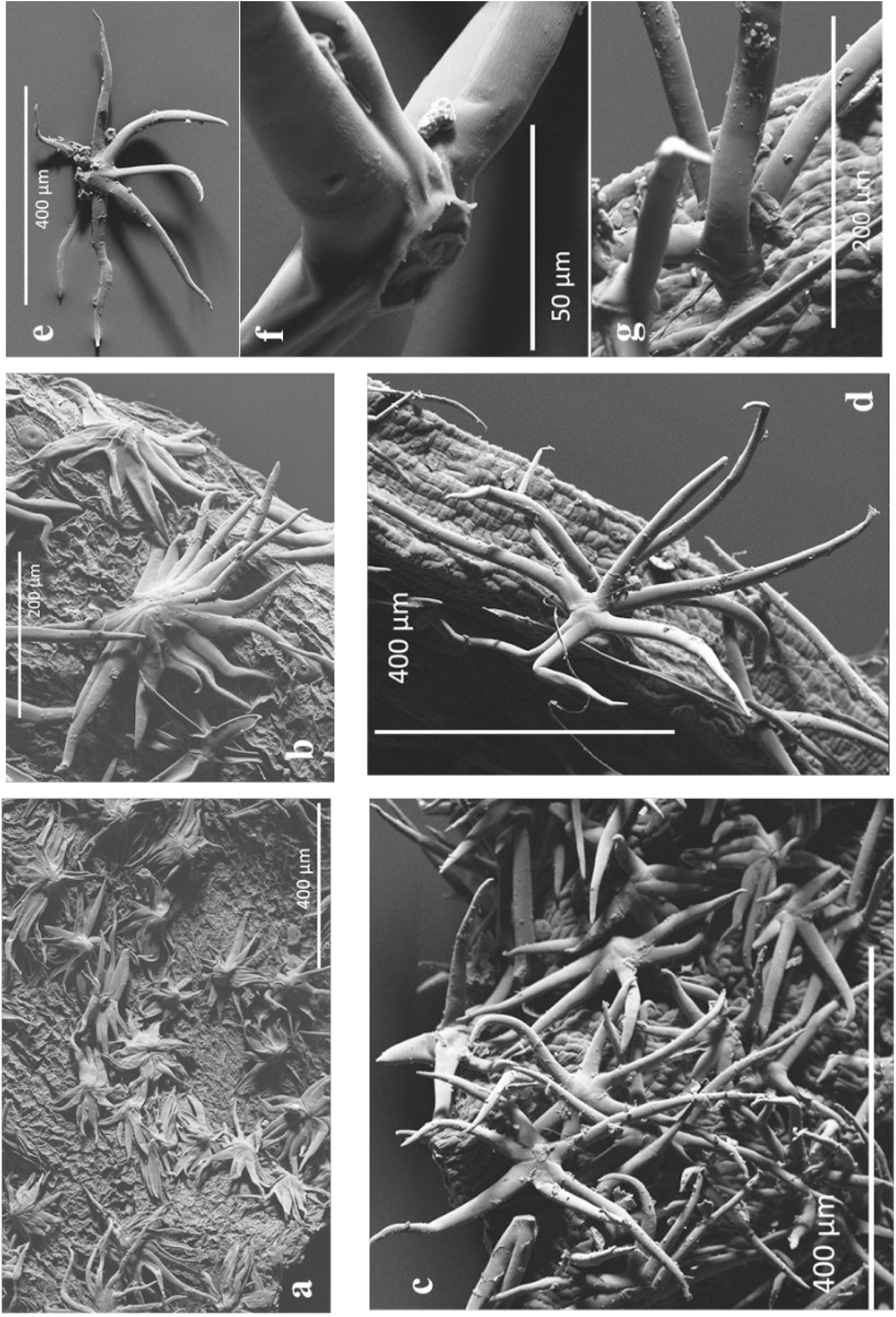
SEM of *Hedera* trichomes: a-b: Spain, Los Barrios (*H. maderensis* subsp. *iberica*); c-g Sicily, Madonie and Alcamo (*H. hibernica*).

Collected plants from Mallorca, Corsica, France and southern Italy have classical *Hedera helix* leaves polymorphism and stellate trichomes.

Trichomes from Sicilian plants seem more polymorphic most of them are stellate type but some are more or less rotate (scale-type related). Unfortunately, most of them fell and remained very scarce in leaves. They have 5-8[10] arms (m = 7.4 ± 1.37); practically footless (Figure 3f), central disc (welding area of arms) often reduced (26µm!), average disc diameter m = 56.6 ± 14.2 µm. The free parts of the different arms of the same trichome are of variable lengths: arm length comprised between 63 and 354 µm (m = 191 ± 81.2) and base width m = 18.7 ± 5.62 µm. Some arms are partly fused (Figure3d) or bifid beyond the central disc.

Plants from Sicily present stellate type trichomes but majorities of them are more or less adpressed (Figure 3c) and then would not match with *Hedera helix* (Ackerfield 2001). Moreover, biometric data match better with the ones of *H. hibernica* than *H. helix*: arms more numerous and shorter (Lum & Maze 1989). Then, all these characters do not allow us to clearly identify species.

### 3.2. Doubts and controversies about genome size

Ivies genomes size presents a good correlation with chromosomes numbers with estimated relationship P_i_ = 0.916 + 0.920 x + 0.077 x^2^ (adjusted R^2^ = 0.995, P<2.2 x 10-16). The relationship estimated from our data is mainly linear with a slightly quadratic component (Figure 4). All parameters depart significantly from zero.

**Figure 4.**
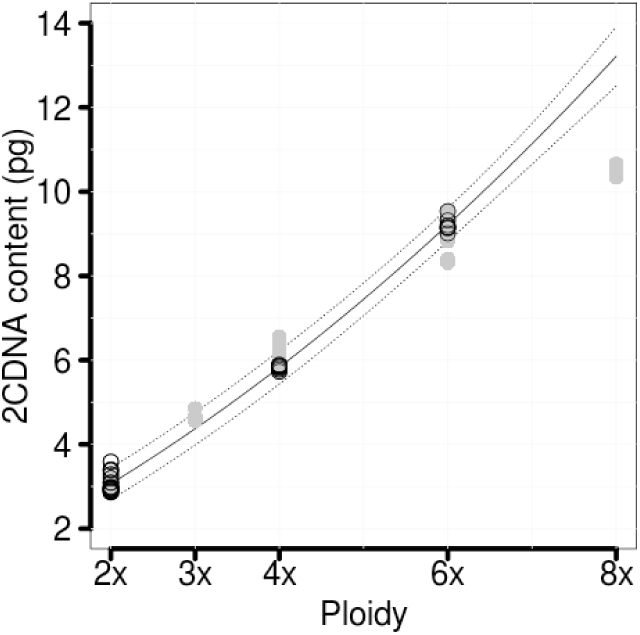
relationship between genome size (2C DNA) and ploidy level in the genus *Hedera*. **Continuous line**: estimated relationship (R2=0.995, P<2.2 10-16) between genome size and ploidy level. **Doted lines**: mean confidence prediction for a probability level equal to 0.95. **Black dots**: 2C DNA values (in picograms) for each plant analyzed. **Grey dots**: genomes size data from Zonnevald et al. (2005) and Green *et al.* (2013).

Our 2C DNA values (Table 2) match well with the data published up to now (Green et al. 2013, see also Figure 4) when considering 2x to 6x range of ploidy. Based on our model, the 8x genome size given by Green et al. (2013) cannot be accurately predicted.

The “*Hedera helix”* 2C value (8.18 pg) given by Marie & Brown (1993) come from an unnamed garden ivy collected in Gif-sur-Yvette (France, Essone), « going to the lab» (S. Brown, personal communication). Then, analyzed leaves belong to one of the many ivies cultivars that are grown abundantly in the surrounding of Paris region. Moreover, internal standard used (*Petunia hybrida*, 2C = 2.85 pg) is not very suitable to measure precisely such large genome. Then this value is not exceptional nor atypical as stated by Obermayer and Greilhuber (2000) but corresponds probably to an hexaploid cultivar (ca 9 pg). Flow cytometry is very suitable to estimate ploidy level in *Hedera* genus as triploid of recent hybridization between *H. helix* and *H. hibernica* have a 2C DNA = ca 4.5 pg (Figure 4). On the other hand, *H. colchica* 2n = 8x = 10.3-10.8 pg (Zonneveld et al. 2005; Green et al. 2013) is lightly different from this pattern, while we would expect a 2C genome size of about 12pg unless plant samples analyzed could be heptaploid hybrids.

## 4. Discussion

Based on trichomes morphology ivies from France, Italy, Corsica and Baleares are *Hedera helix*, DNA amount confirms that they are diploid. Likewise, leaves of Andalusian ivy have an elongated central leaflet, scale-trichomes and plants are hexaploid: they correspond without ambiguity to *H. iberica*.

The dimensions of trichomes vary from an author to another (Lum and Maze 1989; Ackerfield 2001; Valcarcel 2002), which is not surprising due to their great variability on the same branch. Besides in *H. helix* and *H. hibernica* there are many trichomes with intermediate aspects (Valcarcel et al. 2002).

Trichomes morphology does not allow to unambiguously name the plants collected in Madonie and Alcamo forests as one of the three known Sicilian species. In fact, trichomes morphology match better to those of *H. hibernica*: arms shorter, more numerous and adpressed. On the other hand, they are tetraploids. On this point, Sicilian ivy do not correspond to any known Sicilian ivies putatively all diploids: native *H. helix* subsp. *helix*, locally naturalized *Hedera helix* subsp. *poetarum* Nyman or cultivated *Hedera canariensis* (Giardina 2007).

Sicilian ivies leaves look like to *H. helix* ones and most of trichomes are stellate however they are slightly adpressed. Then, if we combine these characters, ivies collected in the northwestern forests belong to *H. hibernica* as the other known tetraploid ivy species (*H. algeriensis*) have scale trichomes (Table 1). While diploids and hexaploids are known throughout *Hedera* genus range, the two tetraploids ivies are confined to the European Atlantic area and Northern Algeria (Figure 1b).

From these observations, it is likely that Sicilian ivies are related to *H. hibernica*. However, we cannot also exclude that it would be a native tetraploid with a great biogeographical interest (cryptic apo endemic).

Moreover, it could well be a hybrid or a naturalized *H. hibernica* that would have been unnoticed on the island. Of the two populations analyzed in Sicily, none corresponds to *H. helix* yet reported as common over a large part of the island.

It seems unlikely that this plant is introduced in Madonie (relatively far from urban centers), but Alcamo forest is largely anthropized. *H. hibernica* has been cultivated for a long time everywhere and has become invasive in various part of northern hemisphere.

In the highly probable case of a past introduction, field studies should be accompanied by examination of the herbarium samples in order to establish the age of the naturalization. In addition, it seems essential to identify again all the ivies previously reported in the literature: obviously several mentions of *H. helix* correspond to putatively *H. hibernica* (taxonomic confusions). It will allow establishing the status of each of these two taxa.

Widely selected by horticulturists and gardeners, ivies have been the subject of a significant trade since at least the beginning of the 18th century. Since these plants are cheap and easy to grow, it is not surprising that they have become invasive. Indeed, it is well demonstrated that selling price and availability on the market are important factors in their dissemination in the field (Dehnen-Schmutz et al. 2007; Pemberton et al. 2009). In North American, ivies began to naturalize on the East coast of the continent in the 1870s and about 60 years later in the West coast quickly becoming invasive with significant ecological consequences on forest dynamics (Clarke et al. 2006; Green et al. 2013; Strelau et al. 2018).

*Hedera* are abundant in various habitats in Europe but difficult to identify, that is why the arrival of allochthonous ivies in natural populations passes unnoticed. In the Mediterranean, the situation appears to be even more complex as native taxa possess all the known levels of ploidy.

On the French Mediterranean coast, *H. helix* is often located, since it requires relatively mesic forests. English ivy (- or Irish ivy - commercial name for several horticultural plants principally selected from *H. hibernica*) is one of the commonest ivies on the market, most sold in urban landscape for decorative purposes but also for soil stabilization and ground cover. Unfortunately, generalized urbanization and development of road infrastructures facilitate the spreading of those cultivated ivies. It allows contact between introduced ivies with the natural stands of *H. helix* so that in certain sectors (near cities and residential areas in particular) the ivies become difficult to characterize (mixture of native species, cultivars and hybrids). This is patent for example in Southeastern France (see Marseille Table2), Liguria, Toscana, Lombardia and Lazio were *H. hibernica* s. l. is well implanted around urban centers. The islands endure the same treatment: in Sicily, ivy cultivars (scale and stellate groups) are widely planted in archaeological sites! The recent expansion of *Xylella fastidiosa* in Corsica, introduced from the United States via horticulturists and intensive agricultural practices (Janse and Obradovic 2010) highlights the massif horticultural imports plants. In Mallorca, the seaside resorts lead to a standardized landscaping and subsequent horticultural plants introduction. If in those two islands *H. helix* remains strongly present in natural areas, in Sicily the ivies of both sampled forests do not correspond to *H. helix*, but to a tetraploid ivy.

These observations are part of a general process in which invasives colonize primarily the coastal areas and low altitudes (especially if they are strongly anthropized), prior to penetrate native habitat. The polyploids (in particular 4x) appearing, in addition, over represented (Verlaque et al. 2002).

In Lebanon, were the only reported native species is *H. helix* (Mouterde 1970); we also noticed various populations of ivies with stellate adpressed trichomes (scale trichomes ivies are largely cultivated as ornamental from lowland up to 1200m). Then it would be necessary to clarify if it corresponds to a particular Mediterranean taxon or if it also belongs to this mix horticultural-invasive-hybrid package.

Replacement of a mesic native species (like *H. helix*) by a cultivar-invasive-hybrid consortium that may develop in drier ecosystems may have some ecological consequences. Rarefaction of wet habitat taxa to the detriment of xerophytics plants is part of a generalized Mediterranean ecosystems transformation (even putatively well-preserved ones like “relict” forest or protected areas). It is all the more devious as it goes unnoticed.

Therefore, fine ivies systematic study is necessary in order to precisely characterize autochthonous species and understand mechanisms of formation and acclimation of those new taxonomic entities in the field. These biological invasions contribute to global changes (change and probable loss of biodiversity…) of which they are powerful indicators. Indeed, if well identified, they appear as valuable markers of current changes.

## Acknowledgements

We thank Spencer Brown (CNRS - Gif sur Yvette) and Nicolas Brouilly (CNRS - Luminy) for their help in flow cytometry and SEM; Fabrice Paranque and André Gilles for English and manuscript revision.

